# Comparative secretomics identifies conserved WAxR motif-containing effectors in rust fungi that suppress cell death in plants

**DOI:** 10.1101/2021.08.18.456800

**Authors:** Rajdeep Jaswal, Himanshu Dubey, Kanti Kiran, Hukam Rawal, Sivasubramanian Rajarammohan, Pramod Prasad, Subhash C Bhardwaj, Humira Sonah, Rupesh Deshmukh, Naveen Gupta, Tilak Raj Sharma

**Author notes:** **Corresponding author** Dr. T R Sharma, Indian Council of Agricultural Research, Division of Crop Science, Krishi Bhavan, New Delhi,110001, India.

## Abstract

Identification of novel effectors with conserved features has always remained a challenge in plant-pathogen interaction studies. The introduction of the genomics era in plant-pathogen studies has led to the identification of significant candidate effectors with novel motifs such as RxLR and dEER motifs. However, in the case of fungal pathogens, limited conserved motifs associated with effectors have been discovered yet. In the present study, we have performed comparative secretome analysis for major plant pathogens of diverse nutrition mechanisms with the aim of dissecting the features underlying their corresponding secretome and conserved motifs. We showed that rust fungi possess the lowest Cell wall degrading enzymes (CWDEs) consortium lower than other biotrophic pathogens. We also showed rust fungi possess the highest secretory superoxide dismutase (SOD) than other studied plant pathogens. Further, we prioritized the candidate secretory effectors proteins (CSEPs) of all the studied pathogens by combining various effector mining parameters to highlight the candidates with potential effector features. A novel WAxR motif in conjugation with the Y/F/WxC (FGC) motif was identified in the effectors of various *P. striiformis* races present globally. The WAxR/WAxR like motifs ( WxxR, WAxx, xAxR) containing effectors were also found in the secretome of other rust fungi. Further, the functional validation of two candidate effectors with WAxR motif from *P. striiformis Yr9* showed that these effectors localize to the nucleus as well as cytoplasm, and are able to suppress BAX induced cell death in *Nicotiana benthamiana*. The mutation analysis of individual residues of the WAxR motif (W, A, R) however did not affect the cell death suppression nor subcellular localization of these effectors. Overall, the current study reports the presence of novel motifs in large numbers of effectors of rust fungi with cell death suppression features.

**Highlights:**

- Secretome analysis of various plant pathogens performed
- A prioritization list for candidate effectors designed
- A novel WAxR motif was discovered in rust fungal effector proteins
- Two WAxR candidate effectors were functionally validated

## 1. Introduction

The plant pathogens follow different nutrition mechanisms explicitly called saprotrophic, necrotrophic, hemibiotrophic, and biotrophic lifestyles. Despite being associated with a different lifestyle, all these pathogens follow a common mechanism for the establishment of infection i.e secretion of small proteinaceous molecules preferably called effectors as these proteins are primarily involved in the manipulation of the host’s defense machinery (Jaswal et al., 2020). To gain entry inside the host cell, these pathogens secrete various cell wall–degrading enzymes (CWDEs) that cause loosening of the cell wall. The successful loosening of the cell wall further causes the transfer of various apoplastic, cytoplasmic effectors as well as virulence determinants that suppress the immunity of the host in numerous ways. Several novel mechanisms have been utilized by pathogens that have been highlighted by recent studies, such as suppressing RNA interference, targeting various cellular organelles of host plants, and maintaining conserved fold in the novel effectors to perform a conserved molecular function(de Guillen et al., 2019; Jaswal et al., 2020; Mukhi et al., 2020; Xu et al., 2019; Yin et al., 2019).

Next-generation sequencing (NGS) technologies have led to the generation of huge genomics and transcriptomic data for plant pathogens. The bioinformatics tools have also been used successfully to identify secretomic differences of the various plant-fungal pathogen using publically available data(Guyon et al., 2014; Jaswal et al., 2019; Ozketen et al., 2020; Sperschneider et al., 2018). However, very few studies have been able to highlight the conserved effector features such as motif or domain in the candidate effector proteins. Despite the huge availability of NGS data, only a limited number of conserved effectors and motifs have been identified and validated to date. The sequence diversity and absence of common features have been a great hurdle for the identification of conserved effector proteins in fungal pathogens. Additionally, the frequent evolution caused by host resistance ( *R* ) genes leading to diversification, moreover species-specific role or functional diversity makes these effectors and their corresponding motifs difficult to identify and develop consensus even using bioinformatics tools (Sonah et al., 2016).

Using computational genomics, two major conserved motifs RxLR and dEER present in several effector members of oomycetes have been identified. These RxLR members constituting the largest effector family in plant pathogens till now are responsible for the translocation of effectors proteins in plant cytoplasm (Dou et al., 2008; Kale et al., 2010; Wang et al., 2021; Whisson et al., 2007). In addition to this, the WY motif and CRN effectors have also been identified in oomycetes. Few other motifs have also been identified however their presence is limited to selected candidates (Jones et al., 2018).

In the case of obligate biotrophs, *Blumeria graminis*, the N terminal Y/F/WxC motif is present in various haustorially expressed secreted proteins (Godfrey et al., 2010). The Y/F/WxC motifs have also been highlighted to be present in haustorially enriched secretory proteins of rust fungi however the exact function of these motifs is yet to be determined(Ozketen et al., 2020; Zhao et al., 2020). In addition to this, a motif sequence [SG]PC[KR]P has been identified in some of the Fusarium secretory proteins, but functional validation of these motifs needs to be done (Ma et al., 2010; Sperschneider et al., 2013). Apart from these, no other conserved motif pattern has been identified in fungal effector proteins using sequence-based studies, especially in rust fungi.

Currently, few universal common features have been explored to mine effector proteins in various studies i) presence of signal peptide, ii) small size ≤300 amino acids and ≥ 3% cysteine content, iii) nuclear localization signal (NLS), iv) repeat sequences, v) signature of positive selection pressure, vi) presence of conserved motif sequences, vii) no conserved domain or pathogenicity related domain reported in other organisms and viii) high expression at the time of infection. Multiple studies have used these parameters to identify candidate effectors across various pathogens (Duplessis et al., 2011; Mesarich et al., 2015; Xu et al., 2020). Apart from these parameters, recently developed tools like EffectorP and Localizer have become useful for prioritizing the effector candidates from total secretomes (Jaswal et al., 2019; Sperschneider et al., 2017, 2016) Although, the common signature features have been frequently explored to identify effector proteins in various pathogens, combining tools like EffectorP and Localizer along with other features can aid in the refinement and better prioritization of these proteins. The hypothesis that candidates that follow the maximum number of parameters, out of the all the candidates analyzed could be prioritized and can be used to mine potential effectors. In the case of rust fungi, hierarchical clustering has been also implemented to find the consensus among candidate effectors(Saunders et al., 2012). Unfortunately not all the proteins follow the selected parameters but probably can be used to increase the confidence score of selecting the protein as a target. Furthermore, expression analysis using publically available data can be utilized as additional support, to target these proteins as the prime candidates.

In the present study, we have followed a comparative genomics approach to analyze the differences among various plant pathogens with varied lifestyles at the secretome level. We used softwares like effectorP and localizer in addition to previously known benchmarks for effector identification and then prioritized the effector candidates by selecting the candidates that followed the highest parameters. We identified a novel candidate effector family with a conserved motif-WAxR among several effector candidate members in *Puccinia striiformis* and other Puccinia species. We further characterized two effector candidates possessing WAxR motif from *Puccinia striiformis Yr9* rust fungi using cell death suppression assay, localization, and site-directed mutagenesis. The present study is the first report of identifying a novel rust-specific effector family with conserved motif in several candidates and having a cell death suppression role in rust fungi.

## 2. Materials and methods

Fungal proteome sequences were downloaded from UniProt and Ensembl database(https://fungi.ensembl.org/index.html). The *P. striiformis* and *Puccinia triticina* proteome were used from the genomic resources generated by Kiran et al.,(Kiran et al., 2017, 2016). The infection of *P. striifromis* was done using a susceptible variety of wheat (Agra Local). The samples were harvested at different time periods 1, 3, 5, 7, 9, and 14 dpi (day post inoculation), immediately frozen in liquid Nitrogen, and further used for RNA extraction. *Nicotiana benthamiana* plants grown up to 3-4 weeks were used for agroinfiltration at 22 °C.

### 2.1 Secretome analysis and functional annotation of fungal secretomes

Secretome analysis was performed using the online secretool pipeline (Cortázar et al., 2014). The non-classically secreted proteins were identified using SecretomeP 2.0 software by using proteome that was not predicted to be classically secreted (http://www.cbs.dtu.dk/services/SecretomeP/). The annotation of secretory proteins was done using NCBI- CD search(Lu et al., 2020). The proteins with no hits were considered as unannotated secretory proteins.

### 2.2 Identification of CWDEs, known effector proteins, and virulence factors across fungal secretomes

Identification of CWDEs was done using the dbCAN database and NCBI CD search (Marchler-Bauer and Bryant, 2004; Yin et al., 2012). The known fungal effector, virulence factor possessing conserved domain was identified by NCBI-CD search. The PHI-base plant interaction database was used to identify pathogenicity genes and virulence factors (Winnenburg et al., 2006).

### 2.3 Phylogenetic, sequence and structure prediction analysis

Phylogenetic analysis of the CSEPs family members identified from *P. striiformis* was done using the maximum likelihood (ML) method. MEGA 7.0 software was used for analysis, models were generated and the best available model was used with 1000 bootstrap runs(Kumar et al., 2016). The sequence alignment was done using MAFFT (https://mafft.cbrc.jp/alignment/server/). Structure prediction analysis was done using PHYRE2 software (Yang et al., 2015)

### 2.4 Orthologous gene identification and selection pressure analysis for common gene clusters

The comparison of orthologous genes in pathogen secretome was done using Orthovenn software (Wang et al., 2015). The genes present in the central cluster were used for the selection pressure analysis. Only one gene was taken from each class as a representative member if there were multiple paralogous genes in one organism in the central cluster. The selection pressure was analyzed using DnaSP 5.0 software (Librado and Rozas, 2009)

### 2.5 Prioritization of CSEPs by combining known benchmark features of effectors

The unannotated secretome of fungal pathogens was screened for potential effector properties using the EffectorP 1.0 (Sperschneider et al., 2016). The secretome was further analyzed for small size (upto300 amino acids) and cysteine content of ≥ 3% using CLC genomics. To find out nuclear localization signal (NLS), and repeat sequences, fungal secretomes were further analyzed by different softwares like Nucpred (Brameier et al., 2007) (NLs identification), localizer (Sperschneider et al., 2017) (localization), xtreme (Newman and Cooper, 2007) and T-rek (Jorda and Kajava, 2009) (repeat identification). To increase the confidence of NL and repeat prediction, common sequences predicted by all four softwares were selected for further analysis. The sequences which contained all the four parameters i.e. high effectorP, small size, NLS and repeat was prioritized 100%, if three parameters followed then 75% and two paramteres then 50%. The analyzed proteins were considered as effector, if followed at least any two parameters (50%).

### 2.6 Motif analysis, amino acid usage and expression analysis of CSEPs

The motif analysis of fungal effector proteins was done by using the MEME suite (Multiple Em for Motif Elicitation)(Bailey et al., 2009). The maximum number of motif identification was restricted to 10. The occurrence of the motif was set to any number of repetitions. The minimum motif width was set to 3 and a maximum of 50. The motif that showed a similar motif pattern was analyzed separately to get a deeper insight into motif similarity. The amino acid usage of the effectors was analyzed by calculating the number of amino acids of the mature fungal effector proteins (without signal peptide). The expression of the prioritized fungal effector was analyzed by using expression data available publically in different studies and using a rust expression browser(Adams et al., 2021).

### 2.7 Identification of WAxR like motif-containing effector candidates in other rusts and plant pathogen secretome

The WAxR effector candidates identified in *P. striiformis Yr9* were used as a query for BLASTp against secretome and predicted proteome of various rust and other plant pathogens analyzed in the study. To further identify the WAxR like effectors in species not analyzed in this study, the BLASTp was also done using conserved region around WAxR motif (10-20 amino acids) as a query against NCBI nr database. The secretome of rust pathogens other than analyzed in the study was also identified as described in section 2.1. The unannotated secretory proteins from the various rust species were analyzed for the WAxR motif using the FIMO tool (https://meme-suite.org/meme/tools/fimo).

### 2.8 Cloning and expression of candidate effectors

Three WAxR motif-containing candidate effectors *Pstr_11677*, *Pstr_09735*, and *Pstr_07126* were selected for cloning based on the prioritization list of the *P. striiformis*. The primers were designed for a full-length coding sequence of the candidate effectors (without signal peptide encoding region). The first-stand cDNA was synthesized using 1 μg RNA isolated from *P. striformis* infection wheat samples from various time intervals 1, 3,5 7, 9, and 14 dpi. The PCR of the candidate effectors using cDNA of various intervals was done. The candidate genes were cloned in pENTR/D/Topo vector, confirmed using PCR restriction digestion and sequencing studies. The confirmed pENTR plasmid clones were used for mobilizing the candidate gene CDS sequence in plant binary vector pGWB408 for overexpression studies and pGWB441 for localization studies using LR clonase II enzyme mix (vector provided by Dr. Tsuyoshi Nakagawa’s lab). The confirmed plasmid clones for overexpression and localization studies were transformed in agrobacterium strain GV3101 and further confirmed by PCR before the agroinfiltration experiment.

### 2.9 Cell death suppression assay, subcellular localization site-directed mutation analysis

The confirmed agrobacterium clones containing overexpression construct were grown at 28 °C. The final cultures were adjusted to an optical density (OD) of 0.4 using MES buffer containing 10 mM 2□(N□morpholino) ethanesulfonic acid (MES), 10 mM MgCl2, pH 5.6, and 150 μM acetosyringone. The experiment was performed in 2-3 leaves of one N. benthamiana plant. The BAX construct suspension was also adjusted to OD 0.4 and infiltrated 24 hrs after candidate genes construct infiltration at the same spots. The pictures were taken after 5-7 days of infiltration. For subcellular localization studies, the agrobacterium culture was adjusted to 0.5 with a resuspension buffer. The images were taken after 48 h at 516 nm using Carl Zeiss Confocal Microscope (Germany). The point mutation in the WAxR motif was done using QuikChange II Site-Directed Mutagenesis Kit per manufacturers’ instruction. The primers were designed using the Agilent mutation primer designing tool. The mutations of the WAxR motif were carried as follows: W (tryptophan) to alanine, A (alanine) to aspartic acid (D), and R (Arginine) into A. The mutated residues containing pENTR clones were confirmed using sequencing and further mobilized into pGWB408 and pGWB441 for overexpression and localization studies respectively.

## 3. Results

### 3.1 Secretome analysis and functional Annotation of fungal Secretomes

Secretome analysis of twenty-six pathogens of different lifestyles revealed contrasting variations in classical secretory protein number. The analysis showed *Monilithophora rorei* had the highest secretome of the total predicted proteome (>800 proteins, 4.97%), the highest among all the studied pathogens, followed by *Magnaporthe oryzae* (640 proteins, 5.05%); however, it was surprising to observe the lowest in *P. triticina* (450, proteins,1.63%) (Fig. 1).

**Fig. 1.**
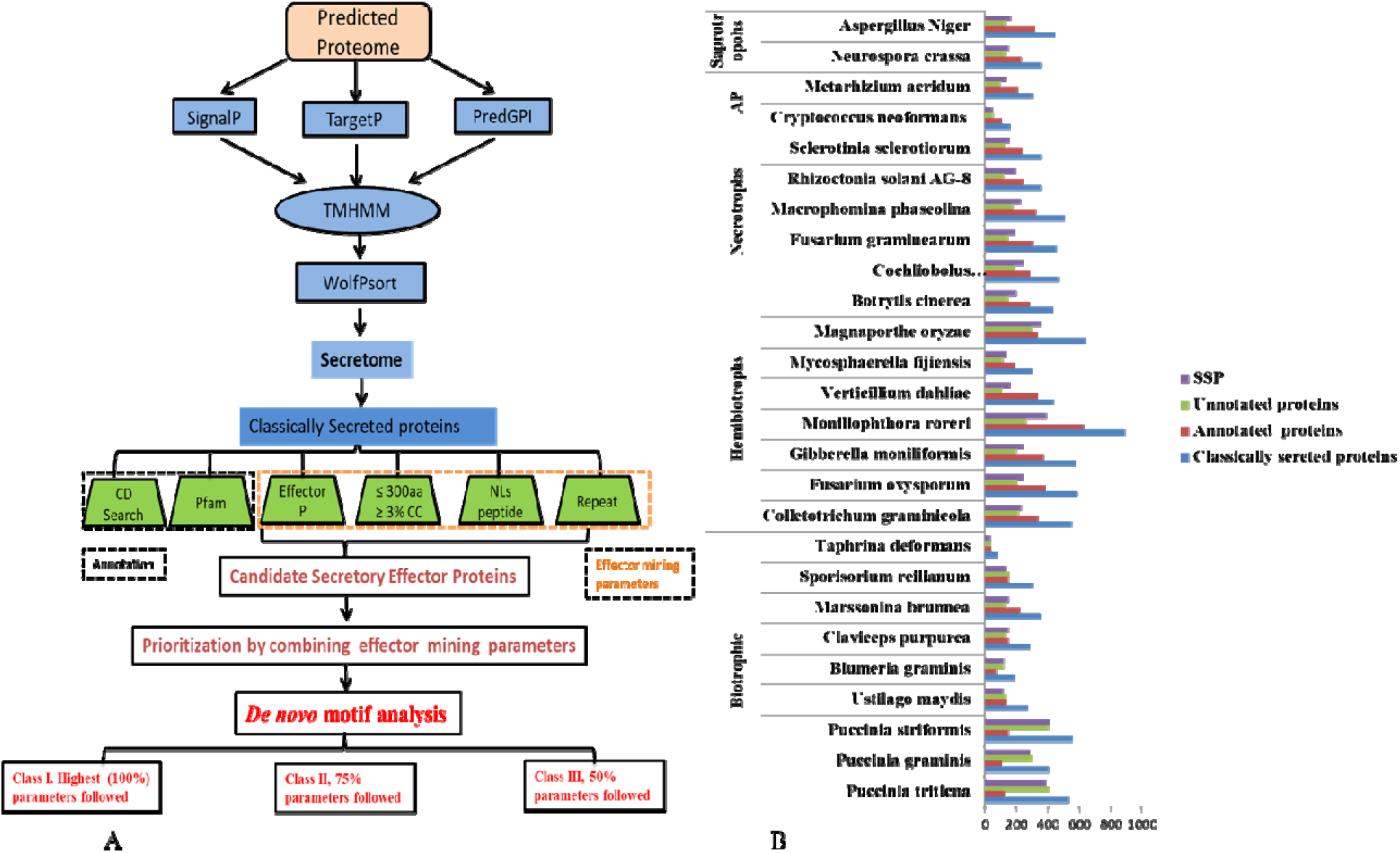
Secretome analysis of plant fungal pathogens. (A) Pipeline designed for the identification of secretome and further candidate effector proteins for various plant pathogens. (B) comparison of Classically secreted proteins, annotated unannotated proteins, and Small secretory proteins (SSPs) of various plant pathogens.

The annotation results of the predicted fungal secretome were consistent with the previous studies and showed that the secretome of obligate biotrophic pathogens contains the highest unannotated proteins. The rust fungi and powdery mildew pathogen were abundant in these proteins and there were 76%, 74%, 73%, and 62 % of unannotated proteins for *P. triticina*, *P. graminis*, *P. striiformis*, and *B. graminis*, respectively. No other pathogens had the secretome that was > 50% uncharacterized (Supplementary Table S1).

### 3.2 Identification of CWDEs, known effector proteins, and virulence factors across fungal secretomes

The identification of CWDEs and known effectors and further comparison was done to find out the differences across different classes of pathogens. The analysis showed the presence/absence of different classes of CWDEs. The lowest no. of CWDEs was found in all the three rust fungi, particularly in *P. striformis* secretome (Figure 2A). It was interesting to note that, even other biotrophic pathogens like *B. graminis*, *C. purpurea*, and *U. maydis* (UM) had higher CWDEs in comparison to the rust fungi (Figure 2A).

**Fig. 2.**
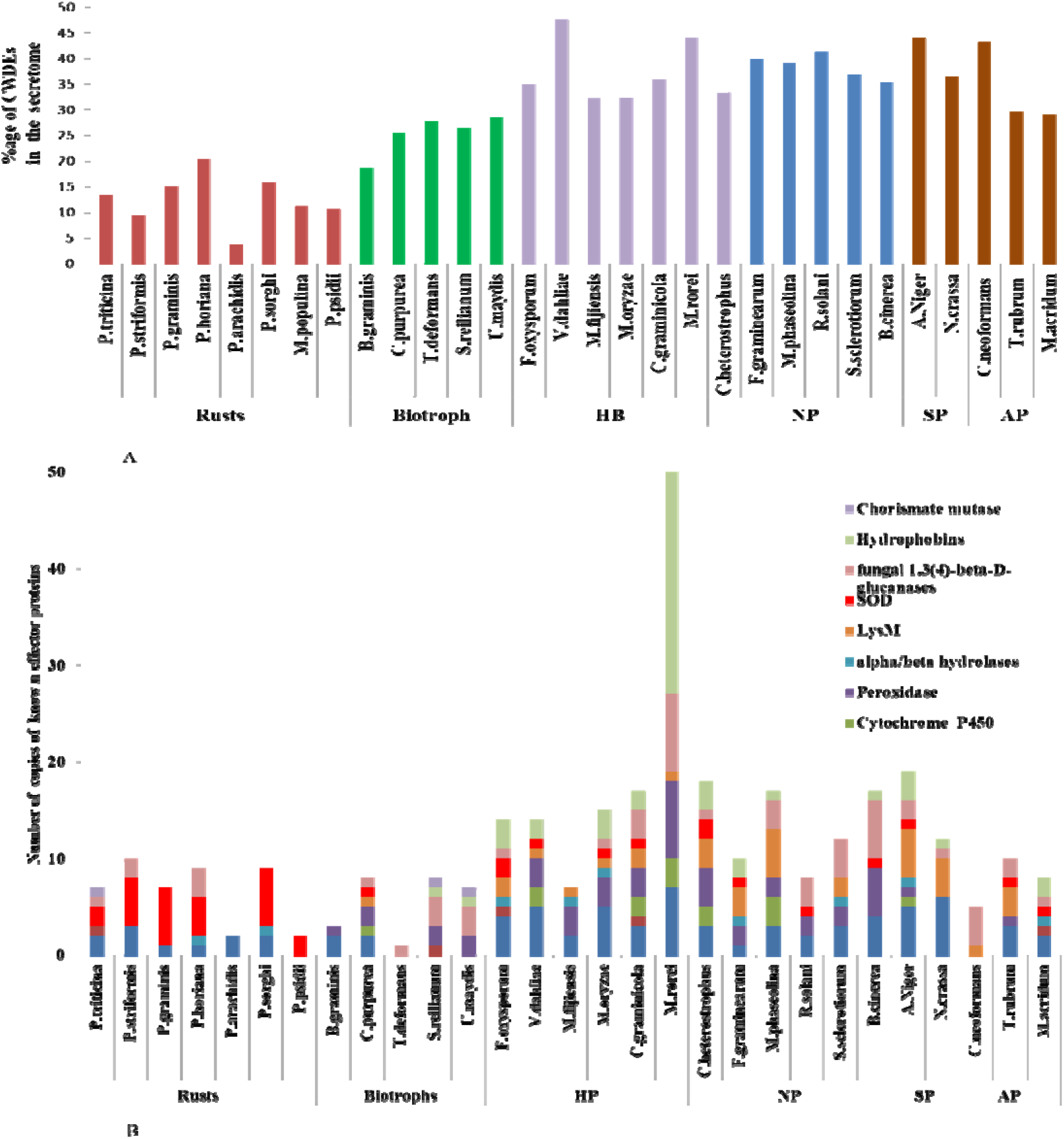
Comparison of CWDEs and known effector and virulence factors in variou fungal species. (A). Number of CWDEs found in the secretome of various plant pathogens. Rust fungi encode the lowest CWDEs in the secretome. (B) The number of known effector proteins present in the various plant pathogens. (HB/HP-hemibiotrophs, NP-necrotrophs SP-saprotrophs, AP-animal pathogens).

In the case of hemibiotrophs, the highest CWDEs were found specifically in *Verticillium dahliae* (VD) and *Monilithphora rorei* (MR) that contain 47.61% and 44.27% of total secretome, respectively. Except for few pathogens in necrotrophs, saprotrophs, and animal pathogens, most of them follow a similar trend to each other in their respective class (Figure 2 A). Various known effectors and virulence factors, like Chorismate mutase (CM), CFEM, FKBP, LysM were also identified in all pathogens. Maximum CFEMs were found in hemibiotrophs in MR (7) (Figure 2 B).

No LysM proteins were found in biotrophs while the highest number was found in *Macrophomina phaseolina* (MR) (5). When mining for SOD, the rust fungi contained the highest number across all pathogens studied. (Figure 2B). In addition to this, fungal hydrophobins were also studied and were present in the range of 0-3 in most of the pathogens but, surprisingly, 24 hydrophobins proteins were present in MR Secretome (Figure 2 B).

### 3.3 Prioritization of identified CSEPs

The parameters mentioned in the materials and methods section (2.5) were combined to prioritize the effector proteins. The analysis identified several effectors in all three major classes of pathogens (Figure 3A). The highest CSEPs were found in *P.striiformis* (218, 43%) followed by *C. purpurea* (109, 37%) and *P. triticina* (161, 30%) (Fig. 3A). Overall, the biotrophs had the highest number of CSEPs and most of them were unannotated. The CSEPs were then classified into three categories: following all the four parameters (100% score), following any three parameters (75% score), or 2 parameters ( 50% score). The candidates with a 100 and 75% score were selected for further analysis. The prioritization analysis results revealed there were 4 proteins in *P. striformis*, 2 proteins in *M. fijensis*, and one protein in *P. triticina*, *C. graminicola*, *M. oryzae*, and *B. graminis* in the 100% score category. There were 77%, 11.8%, 14.3%, 12.7%, 6.2%, 6.7% proteins in *B. graminis*, *P. triticina*, *V. dahliae*, *F. oxysporum C. heterostrophus*, *P. striformis*, and *M.oryzae* respectively that followed 75% score (minimum 3 parameters of CSEPs prioritization). The remaining predicted CSEPs were present in 50% score (two parameters) that followed effectorP score and small size high cysteine content or NLS as a CSEP parameter (Figure 3B).

**Fig. 3.**
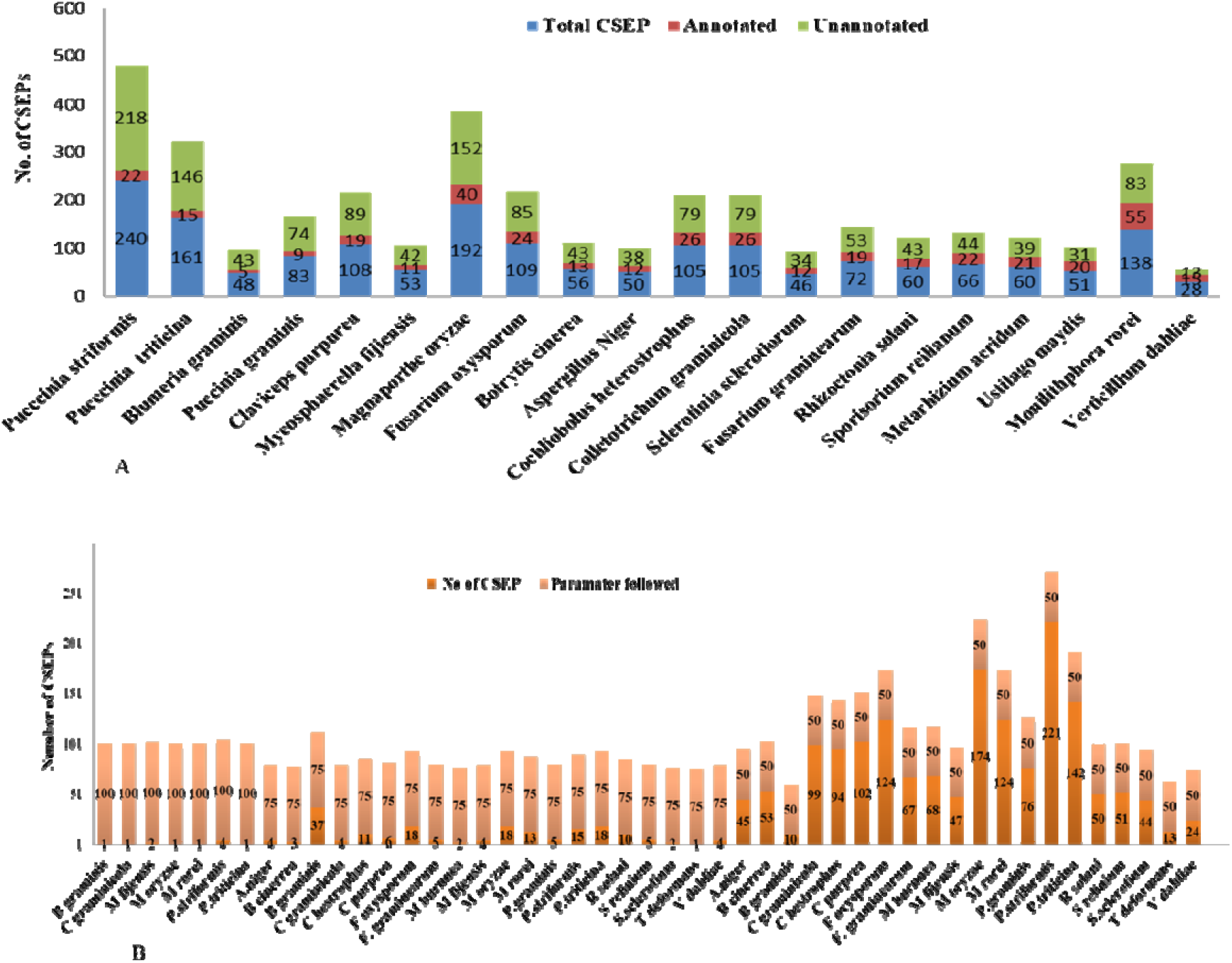
Annotation and prioritization of identified CSEPs identified in various species (A) Annotation of CSEPs for the presence of known conserved domains (B) Categorization of CSEPs of various plant pathogens in three prioritization categories (100% score –for following 4 parameters. 75% score for following 3 parameters and 50% score for following 2 parameters).

*M. burnnea* had a high number of CSEPs in the third category (97.1%) followed by *S.sclerotium* (95.6%) and *C graminicola* (95.2%). Rest all other pathogens except *B.graminis* (9.61%) also had CSEPs in the range of 85-93 % in the 50 % category.

### 3.4 Expression analysis of candidate effectors

The FPKM values (taken from publically available supplementary data from various studies) plotted showed various CSEPs showed high expression suggesting the role of these candidate effector genes in pathogenesis and disease progression. In the case of *P. striformis*, out of the four genes that followed 100% score, one CSEP (*Pstr*_*07126*) showed very high expression in germinating spores stage, and at 7, 9, and 11 dpi (Fig. 4B). The other two genes present in the 75% score category showed high expression *Pstr11677* and *Pstr*_*09735* ( Figure 4 B). In the case of *B. graminis*, out of 77 % CSEP (37CSEP), 15 proteins showed high expression while 7 genes showed moderate expression.at 6,12,24 and 48hpi (Fig. 4C). The expression analysis of CSEPs candidates for *M. oryzae*, *C. graminicola*, *F. graminearum*, and *B. cinerea* also showed that multiple CSEPs encoding genes were high in expression in the infection cycle ( Supplementary Fig S1-S5).

**Fig. 4.**
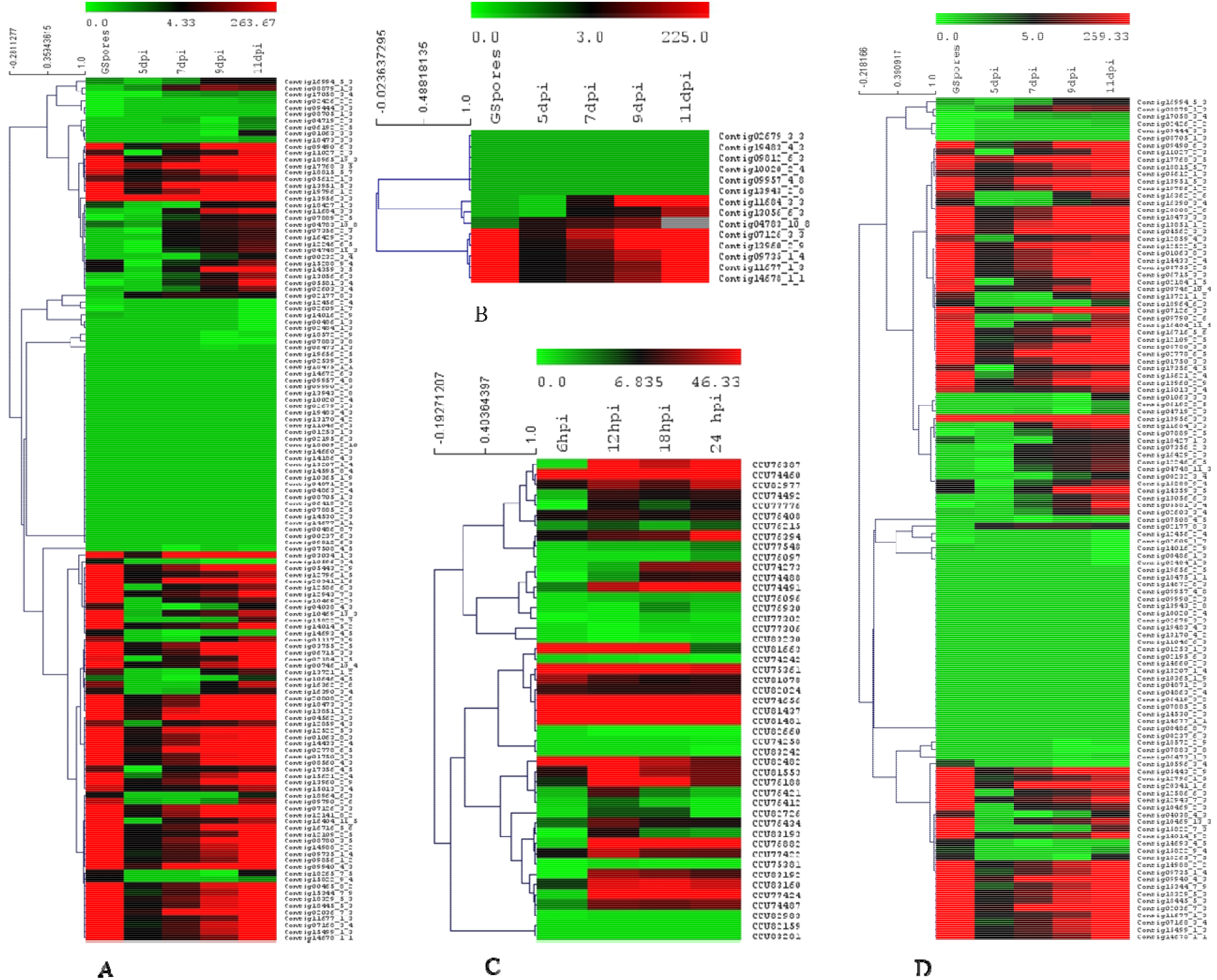
Expression analysis of CSEPs of selected plants fungal pathogens. (A) Expression analysis of CSEPs identified from *P. striiformis* Yr9. (B) Heat map of expression value (FPKM) of the selected CSEPs of *P. striiformis* Yr9 containing WAxR motifs (belonging to 100% and 75 % score categories. (C) Heat map of expression value (FPKM) of the CSEPs *Blumeria graminis*. (D) Heat map of expression values (FPKM) of the CSEPs *Puccinia graminis*.

### 3.5 Sequence and conserved motif repeat and amino acid usage analysis

The conserved motifs were found both at the C-terminal and N-terminal in the case of *P. striiformis (Fig. 5A)* and *Claviceps purpurea*, however, no specific conservation was identified in most of the CSEPs of other fungal pathogens. Among the analyzed CSEP in *C. pupurea*, 3 conserved motifs found in, motif 1, motif 2, and motif 3 were present in 23 proteins covering the full region of all these proteins. (Supplementary Fig. S6 and S7)

**Fig. 5.**
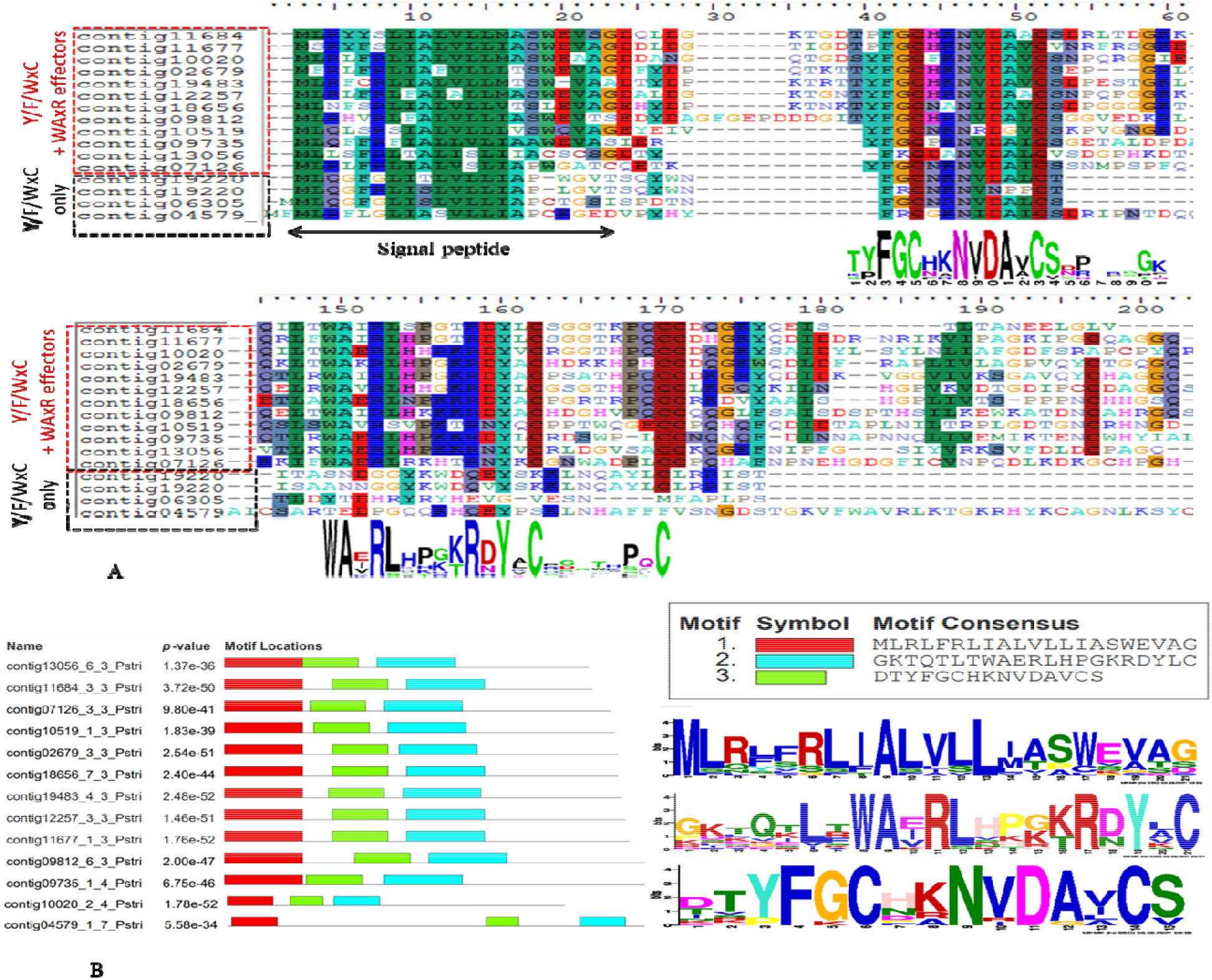
Multiple sequence alignment and motif analysis of WAxR motif effectors from *P. striiformis*. (A) Multiple sequence alignment of effector proteins of *P. striiformis Yr9*. showing Y/F/WxC at N-terminal and WAxR motif at C-terminal. (B). De novo motif analysis identifying WAxR (Cyan) and FGC (green) motif in *P. striiformis* effectors, signal peptide (red).

The multiple sequence alignment of these proteins also shows significant conservation of amino acid residues specifically 6 cysteine conserved residues that were present in most of these proteins (Supplementary Fig. 6). The *P. striifromis* CSEPs had the second-highest motifs of all the pathogens studied. Out of 218 CSEPs given as input, MEME returned 48 proteins with motif prediction. Motif 2 was present in 16 proteins at the N-terminal, after signal peptide. Motif 3 was present in 13 *P. striformis* CSEPs. It was interesting to note that motif 3 was always found associated with motif 2 in 13 proteins. To further gain insight into these motifs, 13 proteins containing these motifs were further subjected to MEME suite and resulted in the identification of three major motifs (Figure 5B). The first motif (red) was the signal peptide region for these proteins. The second motif was the FGC motif, while the third motif was the WAxR motif in the 13 proteins (Figure 5B). The multiple sequence alignment analysis also showed these 13 proteins to significantly conserved residues for the FGC region (Y/F/WXC) followed by the WAxR region (Figure 5A). Out of the four 100% prioritize proteins for *P. striiformis*, Pstr_07126 was the only WAxR motif-containing protein while other WAxR effector proteins (Pstr_11677 and Pstr_09735) were from the 75% score category.

**Fig. 6.**
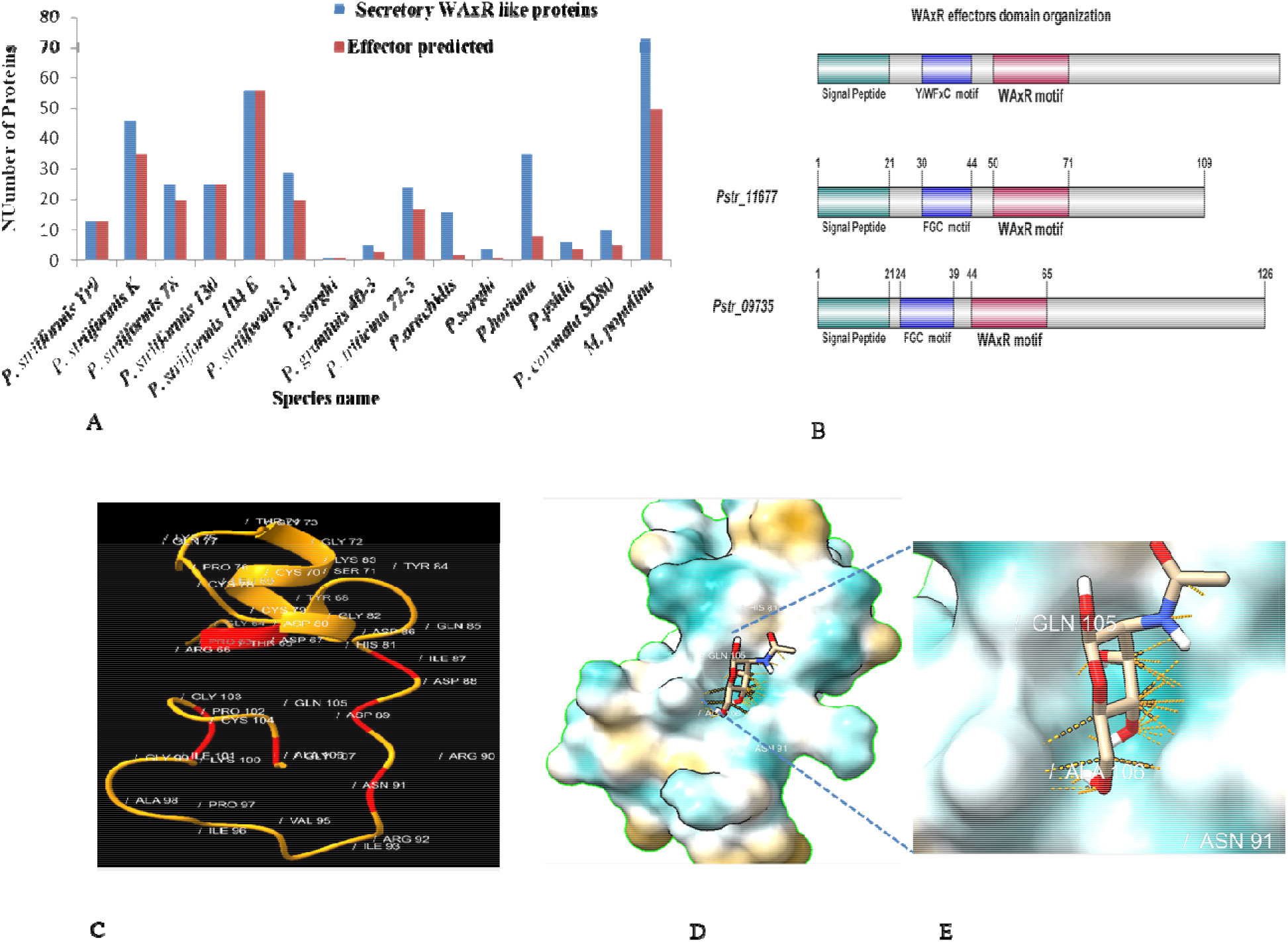
Identification WAxR like effector, domain organization analysis. (A) Identification and comparison of WAxR like effectors in various rust fungi. (B) General domain organization of WAXR effectors in rust fungi. (C) Structural modeling of *Pstr*_*11677* showed similarity to chitin-binding protein. The modeled residues of *Pstr*_*11677*, red area show pocket of chitin-binding residues. (D) and (E) Molecular docking of *Pstr*_*11677* showing binding of chitin (NAG).

The repeat analysis of CSEPs showed that the lowest repeats in CSEP (4.8%-11.25%) were present in rust pathogens and *F. graminearum* particularly in *P. graminis* while the highest was contributed by *V. dahliae* (75%) followed by *S. sclerotium* (58.69%) and *U. maydis* CSEP (55%) (Supplementary Fig. S7). The amino acid usage analysis showed there is no such preference for amino acid usage between effector and non-effector candidates except for cysteine (Supplementary Fig. 8).

### 3.5 Identification of *WAxR*/WAxR like effector candidates in other rusts and plant pathogen genome

The total 5000 BLAST (BLASTp) hits were analyzed against the WAxR motif for plant fungal pathogens other than rust using the NCBI-nr database. Several proteins showed the presence of the WAxR or WAxR like motifs however none of the proteins were secretory nor effector in the analysis except rust fungi. The identification of WAxR like effectors in rusts species apart from analyzed species showed the presence of a large number of WAxR like effectors in their genome. The highest number of WAxR effectors were found in *P. striiformis 104E* (56 proteins) followed by 50 WAxR like candidates in *M. populina*, (Fig. 6 A).

Among all, the *P. sorghi* secretome showed the presence of only one WAxR like effector. Further, the domain organization analysis showed most of these protein possess two major domains. First, the Y/F/WxC motif-containing region that was consistently present at the N-terminal after the predicted signal peptide followed by the C-terminal WAxR motif region in the majority of rust WAxR effectors (Fig. 6B). The C-terminal region after the WAxR motif region was relatively less conserved in these proteins.

The structural modeling of most of the rust WAxR effectors did not reflect any similarity to known fold containing effectors or virulence proteins, however, in the case of *Pstr*_*11677*, we found 26 % sequence similarity to the chitin-binding protein of *Enterococcus faecalis* with modeling confidence of 31.8% (Fig. 6C). The PDB sum analysis also showed similarity to hydrolases with ligand-binding pocket for N-acetylglucosamine (NAG) Further, the molecular docking of *Pstr*_*11677* with chitin and NAG showed significant binding energy −4.8 kcal/mol predicting the putative function of *Pstr*_*11677* (Fig.6 D and E).

### 3.5 Phylogenetic analysis

The phylogenetic analysis of 13 WAxR proteins of *P. striiformis Yr9* clustered together in one group compared to three other proteins containing the FGC motif alone (Figure 7A). To further analyze the group of WAxR proteins in other Indian *P. striiformis* such as Race K, the phylogenetic analysis of WAxR candidates showed clustering in phylogenetic analysis (Figure 7B). The phylogenetic analysis of WAxR and WAxR like effectors identified from various rust fungi classified these into eight groups (Fig. 8).

**Fig. 7.**
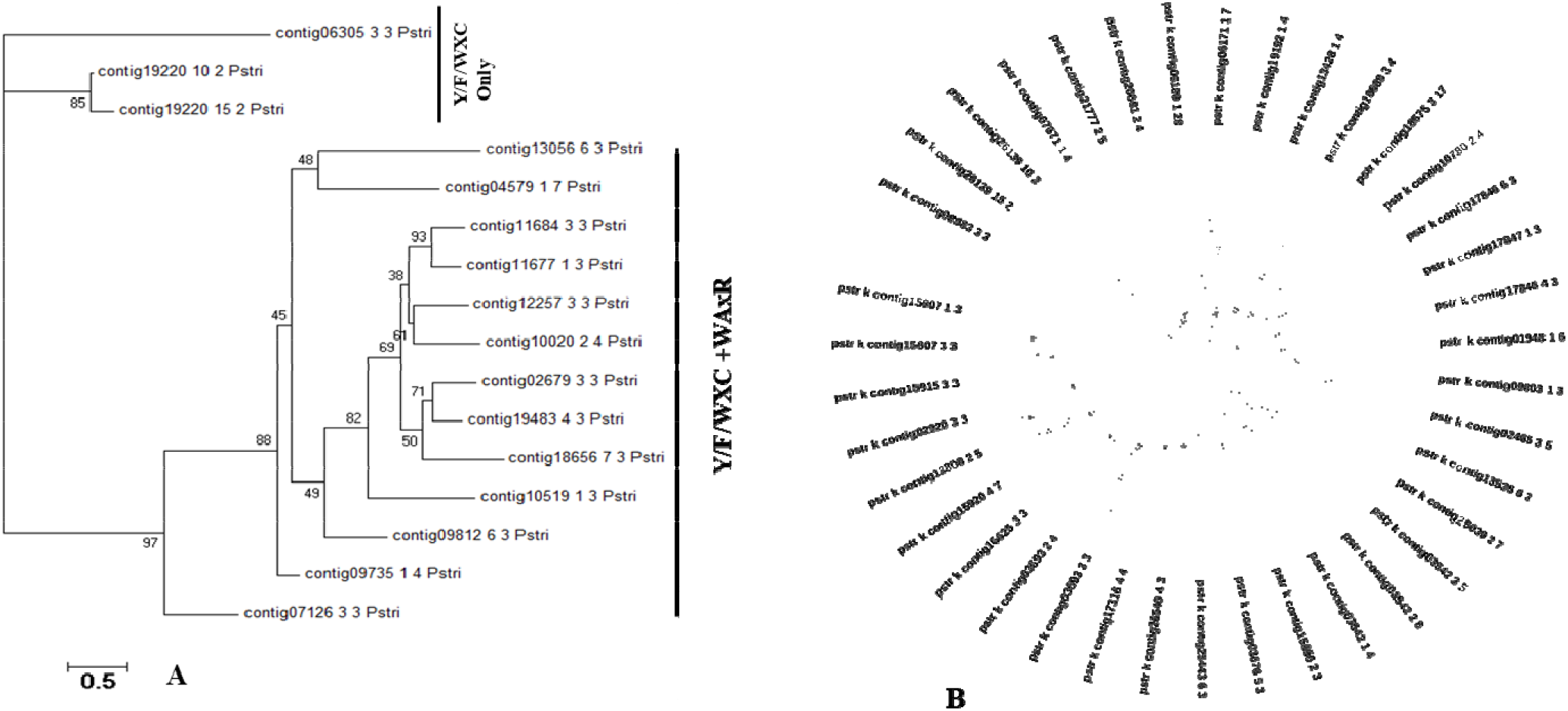
Phylogenetic analysis of candidate effectors of rust fungi. (A) Phylogenetic analysis of WAxR motif-containing effector proteins present in *P. striiformis* Yr9. The WAxR and Y/F/WxC motif clustered separately from the effectors containing Y/F/WxC motif alone (B) Phylogenetic tree of WAxR effector candidates identified from *P. striiformis* Race K.

**Fig. 8.**
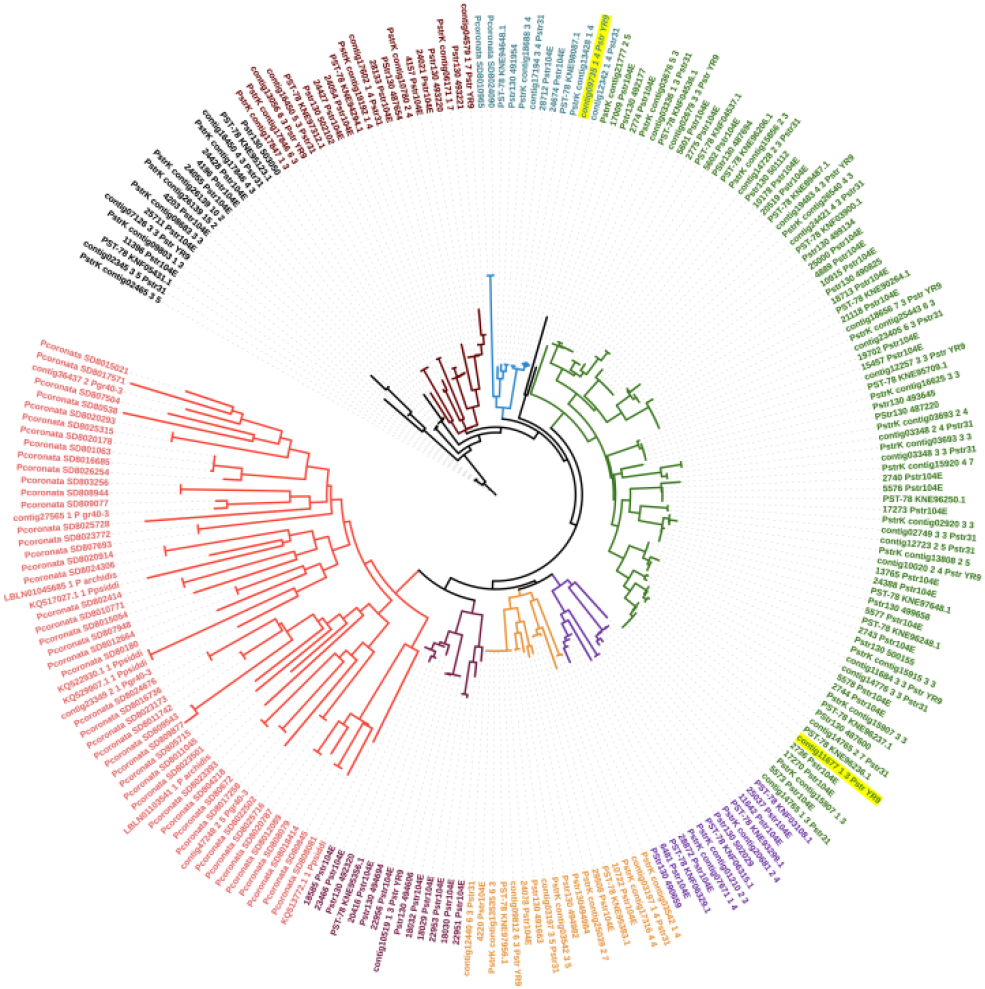
Phylogenetic analysis of candidate effectors of rust fungi. The phylogenetic analysis of all WAxR and WAxR like effectors of rust fungi using the maximum likelihood method (ML) divided these proteins into eight groups (denoted by the different colors). The yellow highlight proteins are the candidates characterized in the present study.

### 3.6. Orthology and selection pressure analysis

The secretome of all fungal pathogens analyzed using orthovenn software revealed most of the genes to be unique, present in singleton for each species. However, few genes were common and present in the central cluster. Three major classes’ necrotrophs, biotrophs, and hemibiotrophs revealed 15, 3, and 13 common genes respectively. Further, the selection pressure analysis of these genes identified 4,1 and 7 genes in necrotrophs, biotrophs, and hemibiotrophs that were on positive selection pressure respectively. The genes that were on selection pressure were mostly glycosyl hydrolase (GH) belonging to class 1, 12, 18, and 61 alpha/beta hydrolases. (Table 1).

**Table 1.**
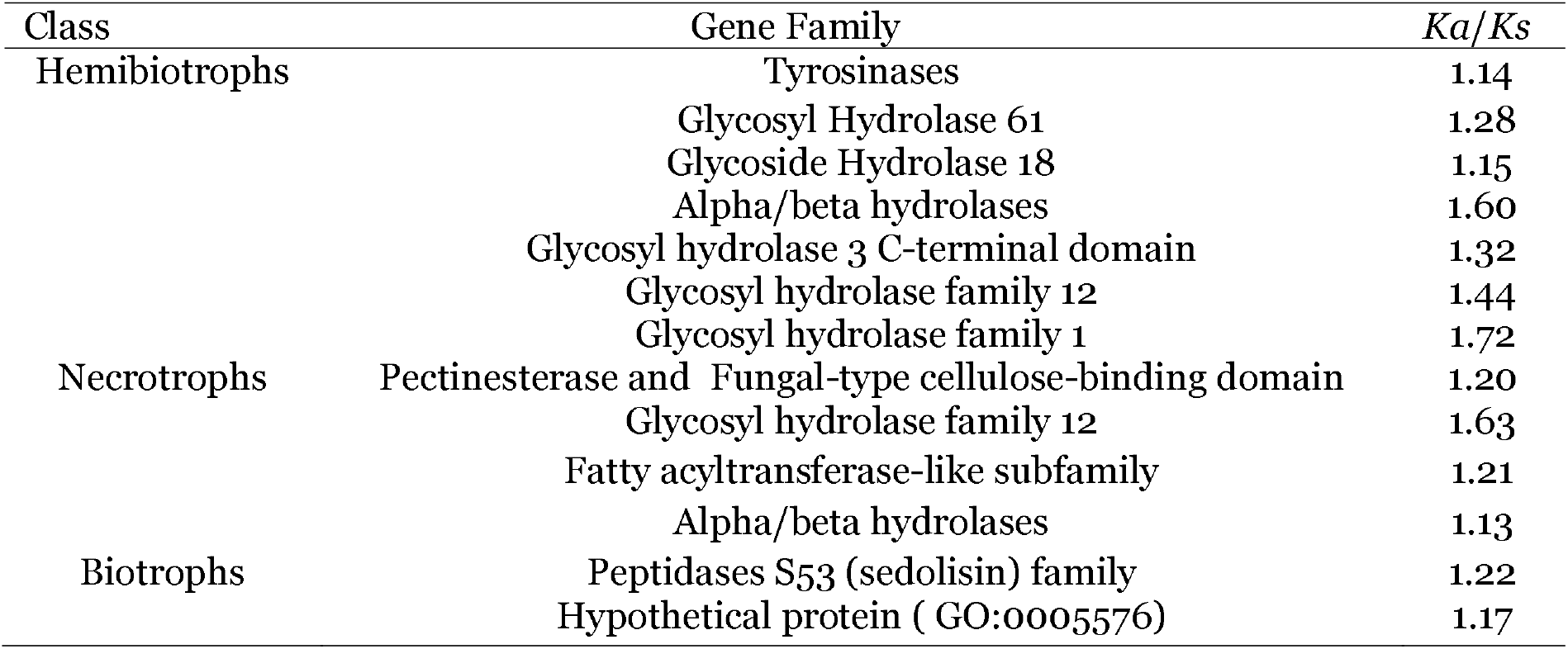
Common proteins found in orthology analysis with positive selection pressure.

### 3.7 Cloning, cell death suppression assay, localization, and mutation of WAxR residues

Out of three genes (*Pstr*_*1726*, *Pstr*_*11677* and *Pstr*_*09735*) two candidates, *Pstr*_*11677* and *Pstr*_*09735* were successfully cloned. The transient overexpression studies of *Pstr*_*11677* and *Pstr*_*09735* studies showed no cell death-inducing phenotype, however, both of these were able to suppress BAX-induced cell death in *N. benthamiana*(Fig. 9A and B). The subcellular localization studies showed *Pstr*_*11677* and *Pstr*_*09735* both localized to the plant nucleus as well as the cytoplasm(Fig. 9C). Further, the mutation of W, A, R residues from the WAxR motif did not affect the cell death suppressing capacity nor the subcellular location of these effectors (Fig. 10).

**Fig. 9.**
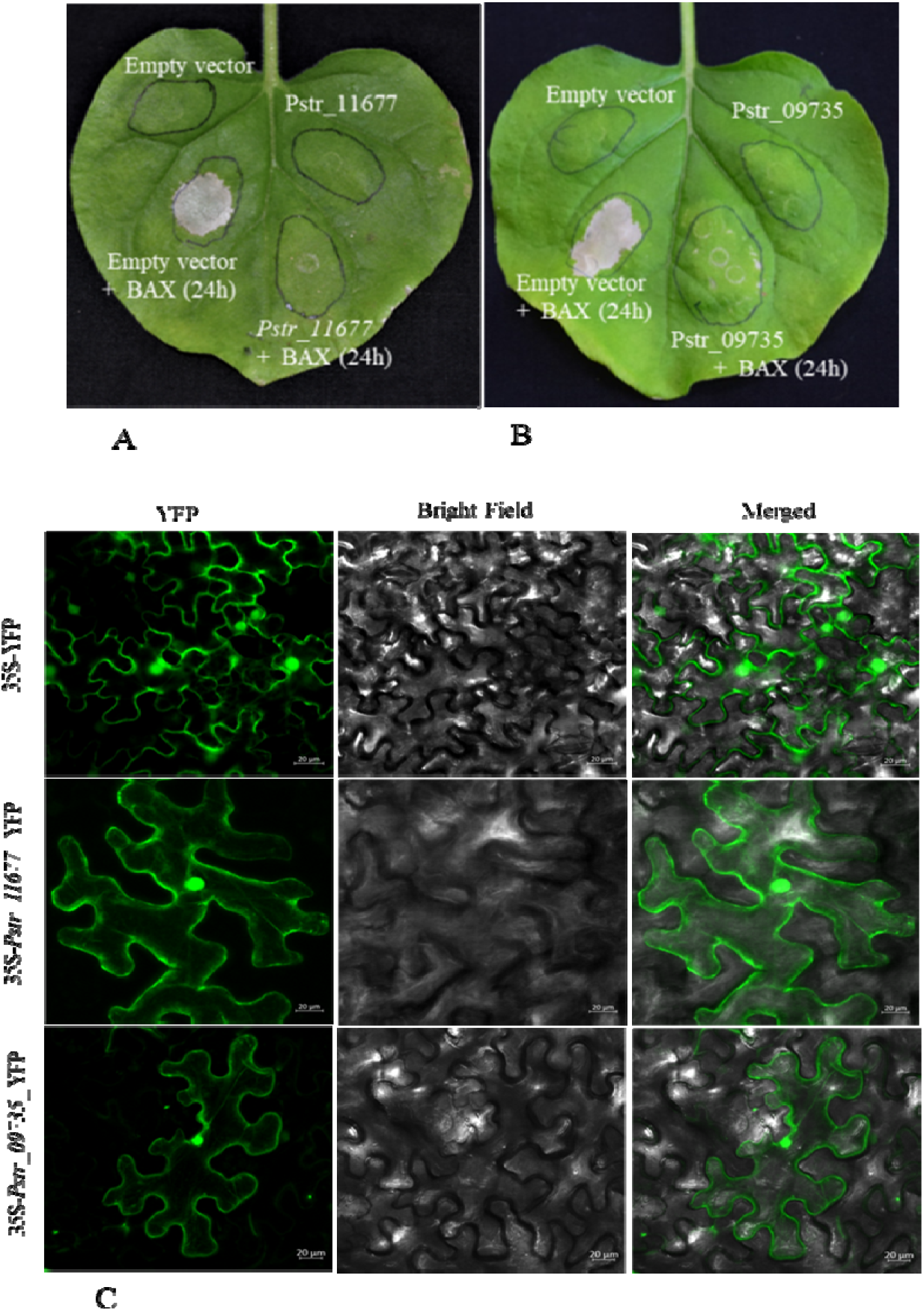
Cell death suppression assay and subcellular localization of WAxR effector. (A) and (B) BAX induced cell death suppression assay using transient agroinfiltration transformation for *Pstr*_*11677* and *Pstr*_*09735* WAxR effector from *P. striiformis Yr9* race. (C) subcellular localization of *Pstr*_*11677* and *Pstr*_*09735* showed these proteins primarily localize to the nucleus and cytoplasm.

**Fig. 10.**
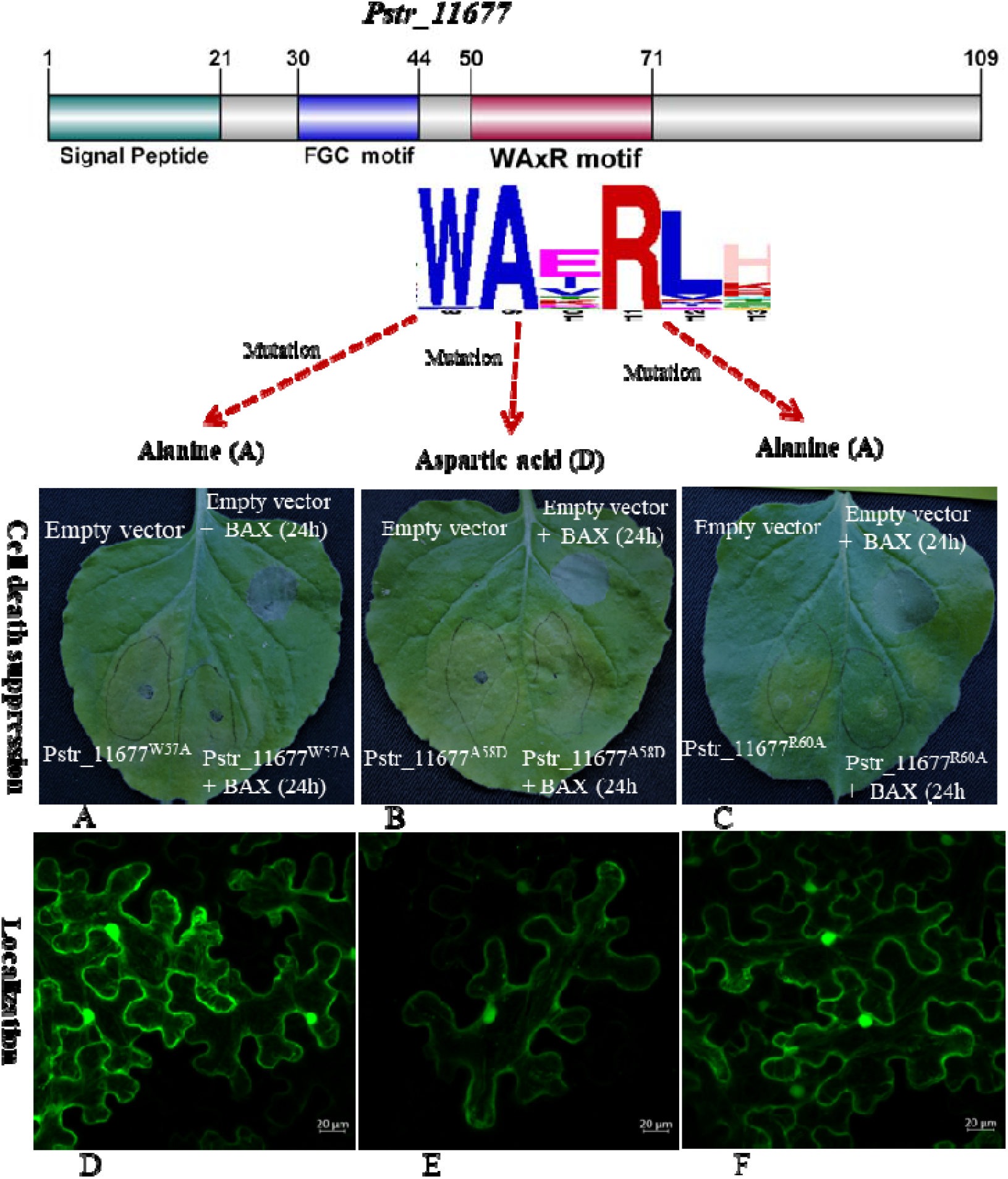
Cell death suppression assay and subcellular localization of WAxR effector after mutations of W, A, R residues. (A) and (D) Cell death suppression and localization of Pstr_1167^W57A^ mutation. (B) and (E) Cell death suppression and localization of Pstr_1167^A58D^ mutation. (C) and (F) Cell death suppression and localization of Pstr_1167^R60A^ mutation.

## 4. Discussion

Secretion of proteins and transferring them to the host cell is one of the primary mechanisms of fungal pathogens to disarm the host. The host as well as pathogens both have a plethora of mechanisms to defeat each other in the war of survival. However, the pathogens have some advantages because of their small life cycle, large spore number, and ability to mutate easily. In addition to this, pathogen develops different mechanism in a unique and faster way to avoid host detection mechanisms than the host that are mostly unknown till now. Moreover, every plant fungal pathogen designs a consortium of secreted proteins that suits its nutritional needs and helps in conquering the host defense. High diversity and specific nutritional choices cause pathogens to have varied secretomes making it difficult to study.

In this study, we have identified, analyzed, and prioritize the CSEPs of various pathogens of different lifestyles. The identified secretome of pathogens describes various features like evolution, infection strategies, mode of nutrition, etc. The secretome analysis showed general trends that pathogens having dual lifestyles have a higher percentage of the secretome in comparison to others pathogens. The top three pathogens *M.oryzae*, *M.rorei*, *C.graminicola* three had the highest secretome of all and belonged to hemibiotrophic (HM) category (Fig.1 B). These results were also consistent with previous analysis (Lowe and Howlett, 2012) (Kim et al., 2016) and further confirmed that high proteome size did not correlate directly to high secretome size. In *U. maydis* only 6522 genes were present, but these genes accounted for 4.18 % of secretome, surprisingly in one of the biotrophic rust fungi *P. triticina* (Kiran et al., 2016) had 32000 genes, however, its secretome accounted for 1.36% of the whole proteome(Fig 1B), but on the other hand, its non-classically secreted proteins was highest of all the pathogens (11480) (Data not shown). Overall, secretory proteins were highest in hemibiotrophs while animal pathogens have the lowest proportion of secretome. The higher secretome in hemibiotrophic pathogens can be supported by the fact that pathogens follow a dual lifecycle where the biotrophic phase requires multiple small secretory proteins that are mostly unknown for restricting the action of the host without causing significant damage and later on require toxins and hydrolases for the killing of the host (Fig.1B and Supplementary Table S1). The biotrophic pathogens especially rust fungal pathogens mostly have high SSPs and unannotated proteins was also reflected in our studies as *P. graminis* and *P. strifromis* has highest SSP followed after *M.oryzae* (Fig. 1B). The CWDEs has also an indispensable role in fungal pathogenesis. Previous studies on CWDEs analysis by showed that necrotrophs and hemibiotrophs have high CWDEs than biotrophs. Similar results were also found in our results, with some exceptions like animal pathogens and saprotroph. *C. neoformans* and *A. niger* had CWDEs almost equal to necrotrophs and hemibiotrophs (Fig. 2A). Although, all the biotrophic organisms contain lower CAZymes when compared with other pathogens, the obligate biotrophs; rust fungal pathogens and powdery mildew showed, even more, less number of CAZymes and high SSP than other biotrophs(Fig. 2A). The results also implied that obligate biotrophic pathogens use less vigorous mechanisms to invade the host and use much more specific ways than other biotrophic pathogens like *U. maydis* and *C. pupurea*. In addition to CWDEs, multiple virulence factors also help in the shaping of the secretome of pathogens. We found a varied number of CFEM effector proteins in all pathogens implying the importance of this effector in pathogenicity (Fig. 2B). Though the function of CFEM effector has not been entirely known till now, it has been known to play functions like signaling, host surface adhesion, and role in infection structure development in multiple pathogens(Liu et al., 2019; Zhao et al., 2020; Zhu et al., 2017). Recently *BcCFEM1*, a Botrytis cinerea CFEM containing proteins is reported to perform multiple functions like stress tolerance, conidia formation, and virulence (Zhu et al., 2017). A recent genome-wide identification has also highlighted a total of 24 CFEM domain-containing proteins however this number containing proteins with the transmembrane region(GONG et al., 2020). Additionally, our analysis showed that CFEM family members are not only restricted to pathogenic fungi but are also present in non-pathogenic fungi like *N.crassa* and even in high copy numbers supporting the role of the protein in the diverse processes (Fig. 2B).

The LysM fungal effectors are, present in multiple plants and animal pathogens and bind with chitin to surpass detection by the host defense system (Dubey et al., 2020; Kombrink and Thomma, 2013; Levin et al., 2017). Surprisingly our analysis revealed that this effector was absent in the secretome of biotrophic plant pathogens except for *C.pupurea*(Zhao et al., 2020) (Fig. 2B). The absence of this effector in these pathogens could be expected with the fact that in *Uromyces viciae-fabae* rust fungi, convert the chitin into chitosan using the chitin deacetylase gene when the mycelium of pathogens penetrates the host stomata (Deising and Siegrist, 1995). The change of chitin into chitosan leads to less affinity of host chitin-binding proteins causing a less vigorous immune response by the host (Deising and Siegrist, 1995). The other explanation could be given that; rust fungal pathogens may bypass this chitin-based detection using some other unknown mechanism.

Among the various defense mechanisms used by the host, the generation of the oxidative burst has proven to be effective as it restricts the growth of pathogens in many ways. The different Reactive Oxygen species (ROS) produced by the host not only kill the pathogen by damaging proteins lipids, cell components but also activates hypersensitive response and play role in signaling pathways leading to activation of a network of defense response (Tenhaken et al., 1995; Torres et al., 2006). In our study, the rust fungal pathogens showed the highest number of SODs than all other pathogens (Fig. 2B). The presence of high SOD proteins in rust pathogens is evident by the fact that damage caused by ROS production causes resistance to host cells against biotrophs in combination with various metabolites, restricting the movement of pathogens to other cells by necrotizing that area. On the other hand, necrotroph pathogens survive these conditions easily by secreting various enzymes and compounds like oxalic acid, catalase, peroxidase, etc. Moreover, necrotroph pathogens flourish easily in areas necrotized by the hypersensitive response, as they take nutrition from the dead cells only. (Able, 2003; Mayer et al., 2001; Rajarammohan, 2021). In addition to this, recently Liu et., 2016 have shown that Zn-SOD of *P. striformis* is secretory and provides increased resistance against oxidative stress induced by the host to *P.striformis* (Liu et al., 2016). Interestingly other biotrophs had no or very less SOD which indicates that rust fungal pathogens use these proteins as a crucial mechanism to tackle the host oxidative defense than other biotrophs. Moreover, we also looked into the status of genes like hydrophobins in the secretome of all pathogens that have reported a wide role in virulence in other pathogens. The hydrophobin proteins have been described to play multiple roles in filamentous pathogens. In *A. fumigatus* these proteins aid in fungal spores to remain undetected from immune response and removal of these proteins by different methods causes spores detection leading to immune system activation (Paris et al., 2003). *Metarhizium brunneum*, also contains multiple hydrophobins genes, differentially express them and play a diverse role in pigment production, conidia production, hydrophobicity, and imparting virulence (Sevim et al., 2012). A hydrophobin gene of *M. grisea* is vital for its development, virulence, and colonization inside the plant (Kim et al., 2005). In our analysis, we found one to three copies of hydrophobin genes in most of the pathogens except *M. rorei* having 24 copies, highest of all studied pathogens (Fig. 2B). The extremely high number of hydrophobins in *M. rorei* showed that hydrophobins could be very important for pathogenesis, so eventually gone substantial expansion in course of evolution.

To look for genes that were common in pathogens and under evolutionary selection pressure, the orthologous genes were analyzed for selection pressure among different pathogens classes. Biotrophic pathogens had a single gene on positive selection pressure annotated as peptidases S53 (sedolisin) a serine peptidase (SP) (Table 1). The SP may be one of the important candidates to target by the host as these have been known to cause direct interaction with the host by degrading the cell wall and pathogenesis-related proteins of multiple hosts and helps in the colonization of pathogens (Orts and Ten Have, 2018; Reichard et al., 2006).

As hemibiotrophic and necrotrophic pathogens are more vigorous in action against the host, the prime barrier that prevents entry of these pathogens inside the plant host is the cell wall. The removal of this barrier requires the expression of various cell wall degrading enzymes like Glycosyl hydrolases (GH) for both classes of pathogens enabling direct contact of these GH with the host (Gruber and Seidl-Seiboth, 2012; Kubicek et al., 2014; Palomares-Rius et al., 2014). Proving this fact, the hemibiotrophic pathogens had seven genes on positive selection while necrotrophs had three, mostly belonging to the GH category (Table 1). The presence of two common genes GH12 and alpha/beta hydrolases in necrotrophs and hemibiotrophs showed these genes may be associated with necrotrophic mode of nutrition for these pathogens. The GH12 of hemibiotrophic oomycete pathogen *Phytophthora* sojae acts as virulence factors having xyloglucanase and β-glucanases activity (Ma et al., 2015). In addition to this, GH12 also triggers pathogen-associated molecular patterns (PAMPs) immunity and causes cell death, when expressed in soybean, *Nicotiana benthamiana*, and other members of the solanaceae family (Ma et al., 2015). In another hemibiotroph fungus *V. dahliae*, two GH12 proteins in association with Carbohydrate-binding module1 also acts as a virulence factor, activating PAMP immunity (PTI) in the host and causes cell death (Gui et al., 2017). Another gene on selection pressure was GH61, now called copper-dependent Auxiliary Activity family 9 (AA9) is a weak cellulose-degrading enzyme that enhances the hydrolysis of lignocellulose when used along with the other lignocellulolytic enzymes (Karnaouri et al., 2014).

The sequence-based periodization process returned various CSEP candidates in the analyzed secretome from the different classes of pathogens (Fig. 3B). Every CSEP candidate followed multiple parameters, so the chances of being an effector also get increased up to some extent. The MEME analysis of CSEPs revealed conserved motifs in some of the biotrophic pathogens like *C. pupurea* and *P. striiformis* (FGC WAxR, P.striiformis) and CTPG, *C. purpurea*) (Fig5 A and S6). Moreover, the phylogenetic analysis of all the 230 WAXR/ WAxR like proteins proved that these effectors are most probably associated with some common families in these pathogens as these clustered in eight groups. Similar to the RxLR effector the WAxR effector could be involved in the diverse defense manipulation process as these proteins show high sequence diversity except at the WAxR motif region.

In biotrophs, very few conserved motifs have been found such as Y/F/WxC and RxLR like motifs in *P.strifromis*, *P.graminis* genes eg. *PS87* till now (Godfrey et al., 2010; Gu et al., 2011; Saunders et al., 2012). No other conserved motifs have been reported in rust CSEPs. The presence of WAxR motif-containing effectors in all major stripe rust races presents worldwide implies these effectors are core candidates for these pathogens. The recently sequenced *P. striiformis* 104E race encoded 56 WAxR motif-containing CSEPs suggesting the expansion of these effectors in *P. striiformis* or specifically in this race. In Indian *P.striformis race K*, *Yr9*, *and 31* we found 35, 13, 20 WAxR effectors respectively suggesting these effectors are significantly present in their secretome. The race 104E has been sequenced with long-read sequencing technologies while other previously sequenced stripe rust genomes including race K, race 78 race 130 are the result of short-read based assemblies, therefore, the WAxR effector members in these races could be higher than current predictions. (Fig. 6A) Apart from *P. striiformis*, the WAxR like effectors were also present in other rusts however comparatively less number except for poplar rust *M. populina* that possesses more than 50 proteins. However, it is worthy to note that these proteins were variants of WAxR effectors. Further functional validation is required whether these suppress cell death similar to WAxR effectors, *Pstr*_*11677* and *Pstr*_*09735* identified from *P. striiformis Yr9* or not. The cell death suppression activity of WAxR effectors suggests that they play a crucial role in host defense manipulation as these effectors were able to suppress BAX-induced cell death (Fig. 7A) To determine, whether the WAxR motif is specifically playing a role in the effector function, the mutation of the individual residues W, A, R for both *Pstr*_*11677* and *Pstr*_*09735* was done however the individual mutations of these residues did not alter its role in cell death suppression or localization. It could be speculated that the complete WAxR motif region might have been involved in the cell death suppression activity as three other residues arginine, tyrosine and cysteines were present downstream of the WAxR motif (Fig. 5)

In addition to the WAxR motif region, the presence of the Y/F/WxC motif in the form of FGC further suggested the function of these proteins in host manipulation. The Y/F/WxC have been previously known to be associate with haustoria forming pathogens specifically rust and powdery mildew (Godfrey et al., 2010; Ozketen et al., 2020; Xu et al., 2020). The transcriptomics of haustoria tissues in different studies has shown the association of Y/F/WxC with candidate effectors exclusively present in haustorial tissue(Ozketen et al., 2020; Xu et al., 2020). Apart from the association with haustorially expressed effector proteins the function of the Y/F/WxC motif remains elusive. Moreover, very few WAxR candidates were devoid of the FGC motif in the study suggesting the crucial function of both the motifs in respect to effector functions and further stating that WAxR effectors could be expressed in haustoria. The expression analysis also showed that one of the functionally validated WAxR effectors *Pstr*_*11677* showed upregulated expression in haustoria compared to germinated spores. The structural modeling of the *Pstr*_*11677* showed similarity to the chitin-binding domain further suggesting the effector function of the WAxR members. Moreover, several studies have reported that most of the N-terminal motifs are involved in the translocation of the effectors similar to the RxLR dEER motif, and C-terminal domain or motif are involved in cell death suppression or induction function (Petre and Kamoun, 2014; Rastogi et al., 2019). Therefore it could be speculated that the FGC motif is involved in the translocation of these effectors from haustoria to plant and the WAxR region could be involved in the effector function (Godfrey et al., 2010). However, the deletion analysis of these two motifs may highlight their specific functions. Overall the identification of the FGC motif-containing WAxR effector family in rust pathogens may provide a baseline and new opportunity to explore the specific function of these effectors in host defense manipulation and utilization of these effectors in durable resistant varieties development.

## 5. Conclusions

The secretomics of plant-pathogen has unraveled the complex infection strategies through the understanding of effector proteins. However, these studies have been largely hampered by the presence of novel effectors in secretome that often lacks conserved features especially in the case of rust fungi. In the present study, using comparative secretomics, we enlighten the effector proteins of plant-pathogen including rusts. We concluded that rust fungi use minimum CWDEs, high SODs, and a novel C-terminal WAxR motif associated with the N-terminal Y/G/WxC motif in several candidate effectors of *P. striiformis* and other rust fungi. The WAxR motif could be a major component of the effector consortium of rust as it was present in several candidate effector proteins having a possible haustorial association due to the presence of the FGC motif. The WAxR effectors also suppress the cell death and localize to the nucleus and cytoplasm proving a crucial component of the defense manipulation system of the host. Further functional analysis of the WAxR effectors in resistance and susceptible host cultivars can elucidate their role as a potential effector or avirulence candidates similar to the RxLR motif-containing effectors.

## Acknowledgments

TRS is thankful to the Department of Science and Technology, Govt. of India, for JC Bose National Fellowship. RJ is thankful to the University Grants Commission (UGC), New Delhi for providing Junior Research Fellowship (JRF).`The authors also gratefully acknowledge Miss Aakriti Mehra for assistance in confocal imaging experiments.

## CRediT authorship contribution statement

**Conceptualization:** Tilak Raj Sharma and Rajdeep Jaswal

**Data curation and lab experiments:** Rajdeep Jaswal

**Data analysis and writing first draft:** Rajdeep Jaswal, Sivasubramanian Rajarammohan, Himanshu Dubey, Kanti Kiran, Hukam Rawal, Humira Sonah, Rupesh Deshmukh, Naveen Gupta

**Funding acquisition:** Tilak Raj Sharma

**Resource:** Subhash C Bhardwaj and Pramod Prasad

**Investigation:** Rajdeep Jaswal

**Methodology and writing final draft:** Tilak Raj Sharma and Rajdeep Jaswal

